# Dissociating activation and integration of discourse referents: evidence from ERPs and oscillations

**DOI:** 10.1101/671933

**Authors:** Cas W. Coopmans, Mante S. Nieuwland

## Abstract

A key challenge in understanding stories and conversations is the comprehension of ‘anaphora’, words that refer back to previously mentioned words or concepts (‘antecedents’). In psycholinguistic theories, anaphor comprehension involves the initial activation of the antecedent and its subsequent integration into the unfolding representation of the narrated event. A recent proposal suggests that these processes draw upon the brain’s recognition memory and language networks, respectively, and may be dissociable in patterns of neural oscillatory synchronization (Nieuwland & Martin, 2017). We addressed this proposal in an electroencephalogram (EEG) study with pre-registered data acquisition and analyses, using event-related potentials (ERPs) and neural oscillations. Dutch participants read two-sentence mini stories containing proper names, which were repeated or new (ease of activation) and coherent or incoherent with the preceding discourse (ease of integration). Repeated names elicited lower N400 and Late Positive Component amplitude than new names, and also an increase in theta-band (4-7 Hz) synchronization, which was largest around 240-450 ms after name onset. Discourse-coherent proper names elicited an increase in gamma-band (60-80 Hz) synchronization compared to discourse-incoherent names. This effect was largest around 690-1000 ms after name onset and was localized to the left frontal cortex. We argue that the initial activation and subsequent discourse-level integration of referents can be dissociated with event-related EEG activity, and are associated with respectively theta- and gamma-band activity. These findings further establish the link between memory and language through neural oscillations.

## 1. Introduction

Discourse comprehension involves the creation of a coherent representation of what a story or conversation is about. A key challenge in this endeavor is the comprehension of ‘anaphora’, words that refer back to previously mentioned words or concepts (‘antecedents’), which are ubiquitous in cohesive text and dialogue (Halliday & Hasan, 1976). Anaphor comprehension is thought to involve the initial activation of the antecedent and its subsequent integration with the discourse representation (e.g., Almor & Nair, 2007; Garnham, 2001; Garrod & Sanford, 1994; Gernsbacher, 1989; McKoon & Ratcliff, 1980; Sanford, Garrod, Lucas, & Henderson, 1983; Sturt, 2003). The neural implementation of these processes is largely unknown, but a recent proposal suggests that they draw upon the brain’s recognition memory and language networks, respectively, and may be dissociable in patterns of neural oscillatory synchronization (Nieuwland & Martin, 2017). To shed further light on this idea, the current electroencephalogram (EEG) study investigated the comprehension of anaphoric proper names embedded in two-sentence mini stories such as “*John and Peter are the best players in the football team. The top scorer of the team was John with thirty goals in total.*” To investigate ease of activation, we manipulated whether the referent in the second sentence (‘John’) had or had not been mentioned before (i.e., was anaphoric or non-anaphoric). To investigate ease of integration, we manipulated whether the referent was semantically coherent or incoherent with the discourse context (top scorer ‘John’ had been introduced as one of the best or worst players). Using a pre-registered protocol for high-resolution EEG data collection, preprocessing and analysis, we examined whether referent activation and integration can be dissociated by event-related potentials (ERPs) and by activity in the theta and gamma frequency bands.

### Referent activation and integration

Psycholinguistic theories of anaphor comprehension recognize the importance of memory representations and processes in forming and maintaining referential relationships (e.g., Garnham 2001; Gernsbacher, 1989; Gerrig & McKoon, 1998; MacDonald & MacWhinney, 1990; McElree, Foraker, & Dyer, 2003; Myers & O’Brien, 1998; Sanford & Garrod, 1989, 2005). Anaphora activate antecedents from the discourse representation, which entails a memory representation of described referents. These activated antecedents are then integrated into the unfolding representation of the narrated event^1^. Activation appears to be a memory-based process that uses relatively superficial information (e.g., lexical and semantic overlap or associations, syntactic gender) as cues to potential antecedents in memory (Gerrig & McKoon, 1998; Martin, 2016; McElree, 2000, 2006; McElree et al., 2003). This involves linking two concepts and recognizing the anaphor as an instantiation of the antecedent despite potential differences in linguistic form (e.g., *book*-*novel*, *John*-*he*). Ease of activation chiefly depends on content overlap of the anaphor with the intended referent relative to other antecedents (McElree, 2000, 2006; McElree et al., 2003). People have difficulty activating a unique antecedent when the content of an anaphor does not distinguish between antecedents (e.g., John in a story about two Johns) or does not match any antecedent (e.g., John in a story about David and Peter), in which case they must introduce a new referent (e.g., Gordon & Hendrick, 1998; Haviland & Clark, 1974; Kennison, Fernandez, & Bowers, 2009; Nieuwland, 2014). Anaphoric reference thus involves a form of recognition memory, i.e. the mnemonic processes for distinguishing old from new information.

Once a referent has been activated, the antecedent representation can be integrated into the unfolding representation of the described event (Almor & Nair, 2007; Garrod & Sanford, 1994; Garrod & Terras, 2000; McKoon & Ratcliff, 1989; Nieuwland & Martin, 2017; Sanford et al., 1983; Sturt, 2003). Semantic integration takes place regardless of whether the expression is anaphoric or non-anaphoric, and is facilitated when the unfolding meaning of the sentence is coherent with respect to the preceding context and consistent with what can be expected based on world knowledge (e.g., Camblin, Gordon, & Swaab, 2007; Graesser, Millis, & Zwaan, 1997; Hagoort, Baggio, & Willems, 2009; Hagoort & Van Berkum, 2007; Kintsch & Van Dijk, 1978; Menenti, Petersson, Scheeringa, & Hagoort, 2008; Nieuwland & Van Berkum, 2006a; Van Berkum, Hagoort, & Brown, 1999; Zwaan & Rapp, 2006).

The theoretical distinction between activation and integration thus suggests a difference not only in timing (i.e., activation precedes integration) but also in the underlying processes (i.e., integration, but not activation, is thought to involve combinatorial processes that compute higher-level meaning from individual word meanings; e.g., Cook & O’Brien, 2014; Kintsch, 1988). Consistent with this distinction in timing, initial activation processes can be observed in earlier reading time measures than subsequent contextual integration processes^2^ (e.g., Garrod & Terras, 2000; Sturt, 2003; see also Staub, 2015). In the current EEG study, we attempted to tease apart the processes underlying activation and integration using ERPs and oscillatory responses, which are complementary electrophysiological measures that provide quantitative and qualitative information about online comprehension (e.g., Bastiaansen, Mazaheri, & Jensen, 2012; Lewis, Wang, & Bastiaansen, 2015).

### ERP correlates of referent activation and integration

ERPs are voltage fluctuations that are associated with a specific event (e.g., onset of a stimulus), and which consist of components whose amplitude, polarity, scalp distribution, latency and/or shape can be used to inform theories of cognitive processing. The ERP components that are most relevant to the current study are the N400 and the Late Positive Component (LPC), which have been associated with memory processing and discourse comprehension.

The N400 component is a negative deflection that peaks between 300-500 ms after the onset of each content word and is largest over centro-parietal electrodes (the difference between two N400 components constitutes the N400 effect; Kutas & Hillyard, 1980, 1984). The N400 is strongly associated with lexico-semantic processing and is often viewed as indexing activation of semantic memory representations (Kutas & Federmeier, 2011). Consistent with this view, the N400 amplitude is smaller for repeated words than for new words (e.g., Petten, Kutas, Kluender, Mitchiner, & McIsaac, 1991), presumably because the lexical or semantic representations of repeated words are already activated by the first exposure. In recognition memory studies, correctly recognized old words elicit smaller N400 amplitude than correctly rejected new words (Curran, 1999; Rugg, Mark, Walla, Schloerscheidt, Birch, & Allan, 1998; Van Strien, Hagenbeek, Stam, Rombouts, & Barkhof, 2005; for a review, see Rugg & Curran, 2007). In studies on sentence and discourse comprehension, repeated referents (e.g., anaphora) elicit smaller N400 amplitude than new referents (Anderson & Holcomb, 2005; Camblin, Ledoux, Boudewyn, Gordon, & Swaab, 2007; Ledoux, Gordon, Camblin, & Swaab, 2007; Streb, Hennighausen, & Rösler, 2004; Van Petten et al., 1991). Notably, the N400 component is not only sensitive to word repetition, but also to the fit between a word and the discourse context. Numerous studies have shown that discourse-coherent words elicited smaller N400 amplitude than discourse-incoherent words^3^ (Camblin et al., 2007; Filik & Leuthold, 2008; Nieuwland & Van Berkum, 2006a; St. George, Mannes, & Hoffinan, 1994; Van Berkum, Brown, & Hagoort, 1999; Van Berkum, Zwitserlood, Hagoort, & Brown, 2003).

In one of the few studies on proper name comprehension, Wang and Yang (2013) found that discourse-coherent proper names elicited smaller N400 amplitude than discourse-incoherent names. Their participants read a two-sentence discourse context that introduced two names along with contrastive characteristics (e.g., *Xiaojin and Xiaochang are both very famous. Xiaojin is a singer, whereas Xiaochang is an actor)*, and a subsequent target sentence containing a proper name that was either coherent or incoherent with respect to this discourse context (e.g., *Yesterday a film producer* [coherent]*/music producer* [incoherent] *came to Xiaochang for collaboration).* The obtained N400 effect suggests that participants associated the names with their characteristics mentioned in the discourse context, and used this information rapidly upon encountering a proper name^4^.

Moreover, several studies suggest that discourse context can override the facilitatory effects of repetition (Camblin, Ledoux, et al., 2007). For example, repeated proper names elicit a larger N400 when their antecedent is highly prominent in the discourse (e.g., *John moved the cabinet because John*…) than when the antecedent is less prominent (e.g., *John and Mary moved the cabinet because John*…; Swaab, Camblin, & Gordon, 2004). This ‘repeated name penalty’ N400 effect may be similar in size to the N400 repetition effect of repeated compared to new names (Camblin, Ledoux, et al., 2007).

It is unclear whether these latter N400 findings reflect the impact of discourse context on referent activation alone or also on referent integration (for discussion, see Almor, Nair, Boiteau, & Vendemia, 2017; Callahan, 2008). That is, discourse-coherent referents maybe be easier to integrate with the context than incoherent referents (Nieuwland & Van Berkum, 2006a; Van Berkum, Hagoort, & Brown, 1999; Van Berkum, Zwitserlood, et al., 2003), or they may be easier to activate (Van Berkum, 2009, 2012), possibly because people predicted their appearance (Almor et al., 2017). Several accounts associate discourse-level integration not with the N400, but with the subsequent LPC component (Burkhardt, 2006, 2007; Brouwer, Fitz, & Hoeks, 2014; Brouwer & Hoeks, 2013; Schumacher, 2011).

The positive-going LPC has a parietal distribution, begins approximately around 500 ms after word onset and can last up to 1000 ms. As with the N400, there is ongoing discussion about the functional significance of the LPC component (for a review, see Van Petten & Luka, 2012). Different manipulations can give rise to LPC effects, and we emphasize that these effects need not index a single cognitive function. Some studies show that the LPC is sensitive to manipulations of sentence- and discourse-level plausibility. For example, words that are semantically related to the sentence context but yield an implausible sentence meaning elicit an LPC effect when compared to plausible control words (also referred to as the ‘semantic P600’; Kim & Osterhout, 2005; Kolk, Chwilla, Van Herten, & Oor, 2003). This effect might be related to the well-known P600 effect obtained for syntactically anomalous or unexpected sentences (but see Leckey & Federmeier, 2019). Based on the association between the LPC component and recognition memory (e.g., Neville, Kutas, Chesney, & Schmidt, 1986; Paller, McCarthy, & Wood, 1988; Van Petten & Senkfor, 1996), Van Petten and Luka (2012) argued that such LPC effects may index retrieval or reactivation of previous words in an attempt to revise one’s earlier parse of the sentence. However, in studies on discourse comprehension, words that introduce new referents have been found to elicit larger LPC amplitude than old words (Burkhardt, 2006, 2007; Schumacher, 2009; Schumacher & Hung, 2012; Wang & Schumacher, 2013; but see Van Petten et al., 1991). Because the amplitude of this LPC effect does not depend on whether the new referent was readily inferable from the discourse context (e.g., ‘the bride’ in a story about a wedding), and thus does not seem to be related to activation processes, it may index the cost of integrating a new referent with the discourse representation^5^ (Brouwer et al., 2014; Brouwer & Hoeks, 2013; Burkhardt, 2006, 2007; Kaan, Dallas, & Barkley, 2007; Schumacher, 2009; Schumacher & Hung, 2012; Wang & Schumacher, 2013).

In sum, both the N400 and LPC appear sensitive to old/new manipulations as well as to discourse-level manipulations, but it is unclear whether processes of referent activation and integration are clearly dissociable using these ERP components. Moreover, ERPs provide only a limited view of neural activity, because they are only suited to investigate event-related activity that is both time-locked and phase-locked to word onset (e.g., Bastiaansen et al., 2012). The current study therefore also employs a time-frequency analysis approach, which can disentangle referent activation and integration via their modulations of neural oscillatory activity.

### Oscillatory correlates of referent activation and integration

Neural oscillatory activity reflects (de)synchronization of neural populations as measured in electrical and magnetic activity (EEG, MEG, ECoG), thought to index transient coupling or uncoupling of functional neural systems (Buzsáki & Draguhn, 2004; Engel, Fries & Singer, 2001; Singer, 2011). The study of neural oscillations offers a window into the functional network dynamics of human cognition that is out of reach from traditional ERP approaches. ERP analysis involves averaging of data segments that are time-locked to stimulus onset (Luck, 2014). This procedure cancels out changes in neural activity that are not phase-locked to stimulus onset (Bastiaansen et al., 2012; Tallon-Baudry & Bertrand, 1999). Stimulus-induced changes in neural activity that are not phase-locked to stimulus onset may therefore be undetectable in ERPs, but can be detected with time-frequency analyses as changes in power within a certain frequency band. In this study, we focus on activity in the theta (4-7 Hz) and gamma (>30 Hz) frequency bands, which have been associated with memory processes and with semantic aspects of language comprehension.

Only a handful of studies have applied time-frequency analysis to investigate anaphor comprehension (Boudewyn et al., 2015; Heine, Tamm, Hofmann, Bösel, & Jacobs, 2006; Meyer, Grigutsch, Schmuck, Gaston, & Friederici, 2015; Nieuwland & Martin, 2017; Van Berkum, Zwitserlood, Bastiaansen, Brown, & Hagoort, 2004). In a recent study, Nieuwland and Martin (2017) reported time-frequency analyses of four EEG datasets that compared referentially ambiguous with unambiguous, coherent anaphora (Martin, Nieuwland, & Carreiras, 2012; Nieuwland, 2014; Nieuwland, Otten, & Van Berkum, 2007; Nieuwland & Van Berkum, 2006b), and that each had demonstrated an Nref effect (i.e., enhanced frontal negativity for ambiguous compared to unambiguous anaphora^6^) in the ERP waveforms. In the time-frequency analyses, referentially coherent anaphora elicited an increase in 35-45 Hz (low gamma) power around 400-600 ms and an increase in 60-80 Hz (high gamma) power around 500-1000 ms. The low and high gamma effects were localized to the left posterior parietal cortex and the left inferior frontal-temporal cortex, respectively. In conjunction with these source localization results and previous literature, Nieuwland and Martin (2017) interpreted the low gamma effect as reflecting referent activation by the brain’s recognition memory network (Gonzalez et al., 2015; Wagner, Shannon, Kahn, & Buckner, 2005), while the high gamma effect was taken to reflect integration with the sentence context (Bastiaansen & Hagoort, 2015; Fedorenko et al., 2016; Peña & Melloni, 2012).

Nieuwland and Martin (2017) did not observe modulations in the theta range, but suggested that theta effects may occur in a comparison of old and new referents. In studies on recognition memory, correctly recognized old items elicit enhanced power in the theta band compared to correctly rejected new items (Burgess & Gruzelier, 1997, 2000; Klimesch, Doppelmayr, Schimke, & Ripper, 1997; Osipova, Takashima, Oostenveld, Fernández, Maris, & Jensen, 2006; Van Strien, Hagenbeek, Stam, Rombouts, & Barkhof, 2005; Van Strien, Verkoeijen, Van der Meer, & Franken, 2007; for a review, see Nyhus & Curran, 2010). Theta-band activity has therefore been linked to the process of matching a current item to memory representations of previous items (Chen & Caplan, 2016; Jacobs, Hwang, Curran, & Kahana, 2006) and may originate from the hippocampus and related memory structures (Bastiaansen & Hagoort, 2006; Klimesch, Doppelmayr, Schwaiger, Winkler, & Gruber, 2000; Nyhus & Curran, 2010). This memory matching process involves reactivation of a previously encountered item, even when no explicit recognition judgement is required.

In language comprehension research, theta-band activity has been associated with retrieving lexical-semantic information from long-term memory (Bakker, Takashima, Van Hell, Janzen, & McQueen, 2015; Bakker-Marshall, Takashima, Schoffelen, & Van Hell, 2018; Bastiaansen, Oostenveld, Jensen, & Hagoort, 2008; Bastiaansen, Van der Linden, Ter Keurs, Dijkstra, & Hagoort, 2005; Piai et al., 2016; Strauß, Kotz, Scharinger, & Obleser, 2014). It is unknown whether theta activity also plays a role in referent activation, but we consider this a reasonable prediction from the memory-based view of anaphor comprehension (e.g., Martin, 2016; Nieuwland & Martin, 2017; Sanford & Garrod, 2005). Here, we sought support for the role of memory activation in anaphor comprehension, and we tested whether anaphoric (repeated) referents increase oscillatory activity in both the theta band and the low gamma band compared to non-anaphoric (new) referents. In addition, we sought further support for the role of high gamma activity in integrating referents with the unfolding discourse representation (Nieuwland & Martin, 2017).

### The present study

Our participants read two-sentence mini stories containing proper names (see Table 1 for an example). We opted to use proper names because they contain less semantic content than noun phrases, minimizing the impact of long-term semantic memory representations on the obtained results (e.g., Semenza, 2006; Semenza & Zettin, 1989). Each story introduced two entities in the context sentence, and mentioned one referent in the target sentence. We investigated referent activation and integration by orthogonally manipulating ease of activation (whether the referent was repeated/old or new) and ease of integration (whether the referent rendered the target sentence coherent or incoherent with the preceding discourse).

**Table 1.**
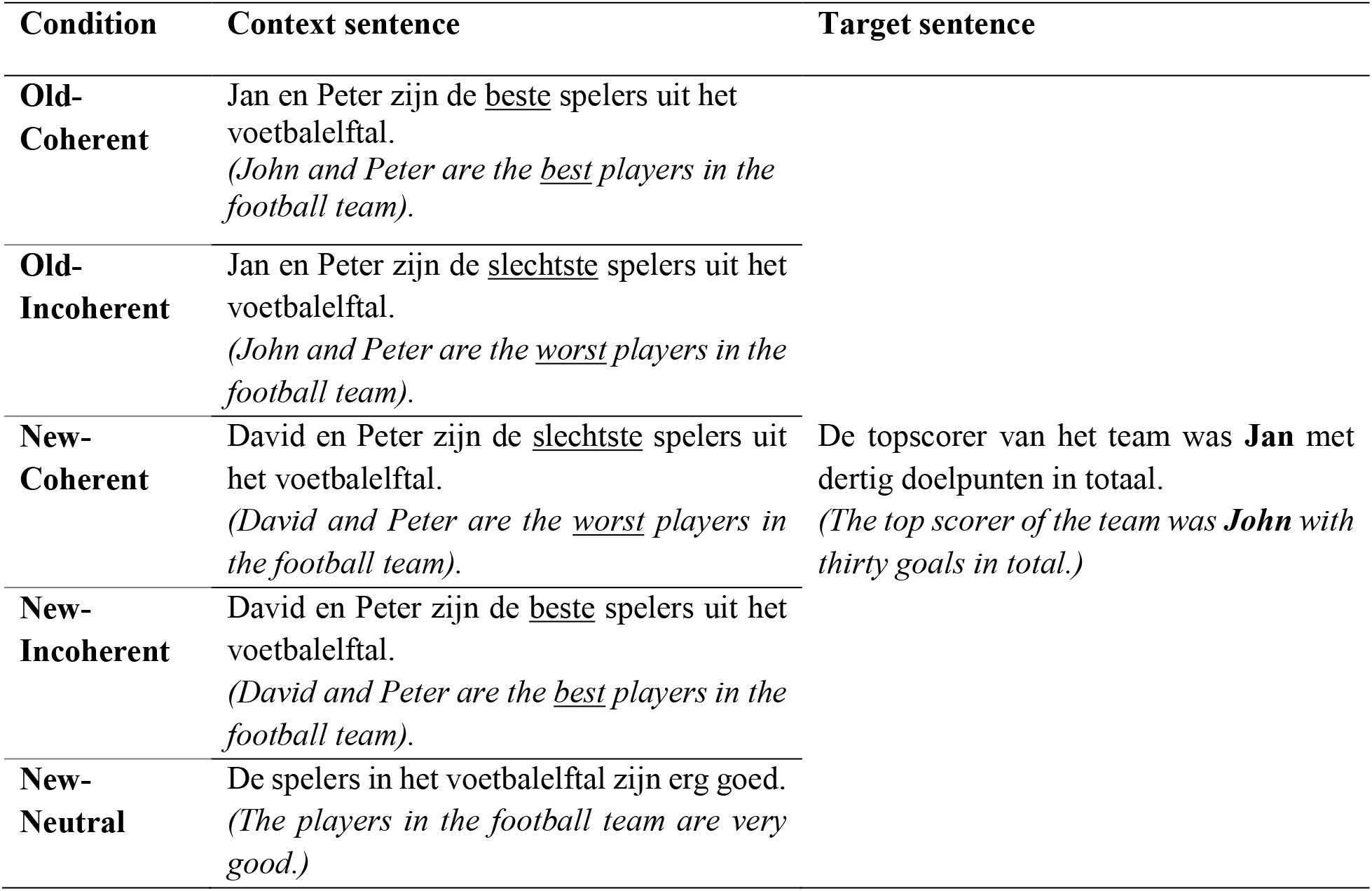
Example stimulus item, containing all five conditions of an original Dutch two-sentence mini story along with an approximate English translation. For expository purposes only, words that differ between the coherent and incoherent conditions are underlined, and the critical proper name is printed in bold. The full set of materials is available on https://osf.io/nbjfm/.

In addition, we included a ‘new-neutral’ condition in which the context did not mention specific names but described a group of people (e.g., *the players in the football team*), thereby providing a non-specific antecedent. This condition allowed us to establish whether the brain’s response to new referents genuinely reflects the introduction of a new referent or perhaps just the mismatch with given antecedents.

Using a pre-registered protocol for high-resolution EEG data collection, preprocessing and analysis, we examined whether referent activation and integration are dissociable by ERPs and/or by oscillatory activity in the theta and gamma frequency bands. We restricted our pre-registered ERP analyses to the N400 and LPC components, and our time-frequency analyses to the theta (4-7 Hz), low gamma (35-45 Hz) and high gamma (60-80 Hz) frequency bands.

We hypothesized that old referents would be easier to activate than new referents, and that this difference would reveal itself in N400 amplitude, with old referents eliciting lower N400 amplitude than new referents. We further considered possible effects of discourse coherence on N400 amplitude. If coherent referents elicit lower N400 amplitude than incoherent referents regardless of whether the referents are old or new, this would suggest that coherent referents are easier to integrate than incoherent ones, supporting an ease-of-integration effect on N400 activity (Brown & Hagoort, 1993; Hagoort et al., 2009). Instead, if old-coherent referents elicit lower N400 amplitude than old-incoherent referents but no equivalent effect is observed for new referents, this would support an ease-of-activation effect on N400 activity (Kutas & Federmeier, 2000; Lau, Almeida, Hines, & Poeppel, 2009; Van Berkum, 2009, 2012). That is, whereas context may facilitate activation of an old-coherent referent (by activating the referent before it appears), it cannot facilitate activation of a new and unpredictable referent.

Alternatively, it is possible that N400 activity only reflects referent activation whereas the subsequent LPC component reveals an integration effect. Such an effect could reflect processes initiated by both repetition and coherence. That is, new referents might elicit larger LPC amplitude than old referents, reflecting the cost of integrating a new referent with the discourse representation (Brouwer et al., 2014; Brouwer & Hoeks, 2013; Burkhardt, 2006, 2007; Kaan et al., 2007; Schumacher, 2009; Schumacher & Hung, 2012; Wang & Schumacher, 2013). In addition, LPC amplitude may be smaller for coherent than for incoherent referents, reflecting facilitated discourse integration for coherence referents (similar to ‘semantic P600’ effects; Kim & Osterhout, 2005; Kolk et al., 2003).

For the time-frequency domain, we hypothesized that effects of referent activation and integration would be observed in different time windows and different frequency bands. We predicted an increase in theta-band (0-1000 ms) and low gamma-band (400-600 ms) power for old compared to new referents (Nieuwland & Martin, 2017). For a later time window (500-1000 ms), we predicted an increase in high gamma-band power for coherent compared to incoherent referents, which may originate from left inferior frontal-temporal regions (Nieuwland & Martin, 2017). As with the ERP patterns, we considered the possibility that the effect of coherence differs for old and new referents.

Although not germane to our main hypotheses, the new-neutral condition enabled us to test whether the process of activating a new referent depends on whether the context already contained specific referents or not. Because the neutral context contained an unspecified set of referents, we hypothesized that a new referent in this condition would incur referential uncertainty and elicit a sustained, frontal negative shift (Nref effect) when compared to the new-coherent and old-coherent conditions.

## 2. Methods

### Preregistration

The design and settings of our analysis procedures (i.e., preprocessing, time-frequency analysis, and statistical analysis) have been preregistered at Open Science Framework (https://osf.io/nbjfm/). Non-preregistered analyses are designated as exploratory.

### Participants

We pre-registered a sample size of 40 participants after exclusions. In total, 45 native speakers of Dutch participated in the experiment for monetary compensation. They had normal or corrected-to-normal vision, were not dyslexic, and were right-handed. After receiving information about the experimental procedure, participants gave written informed consent to take part in the experiment, which was approved by the Ethics Committee for Behavioural Research of the Social Sciences Faculty at Radboud University Nijmegen in compliance with the Declaration of Helsinki. We excluded 3 participants who reported left-handedness after the experiment. We excluded 2 participants from the ERP analysis due to a low number of trials after artifact rejection, and excluded 1 participant from the time-frequency analysis for that reason. Therefore, the ERP results are based on 40 participants (30 females, mean age = 23 years, range = 19-33 years), and the time-frequency results are based on 41 participants (29 females, mean age = 23 years, range = 19-33 years).

### Experimental items

We created a total of 225 two-sentence mini stories^7^. The first sentence of each mini story varied over the conditions, but the second, target sentence was identical for all conditions. The first sentence introduced two people using conjoined proper names (e.g., *John and Peter*). This type of embedding reduces the prominence of both referents, making subsequent use of a repeated name more felicitous than when only one proper name has been introduced (Albrecht & Clifton, 1998; Almor et al., 2017; Ledoux et al., 2007; Swaab et al., 2004). The second half of the first sentence added characteristic information about these two people. This information was manipulated using antonymic expressions regarding social or personality characteristics (e.g., being friendly/unfriendly), physical characteristics (e.g., being strong/weak) or behavioral characteristics (e.g., getting good/bad grades). In addition, the first sentence always described a reference group to which both referents belong (e.g., *the players in the football team*). We added this group to suggest the presence of yet other referents, thereby facilitating the introduction of a new referent. The second sentence described an action or event involving a proper name (the critical word; CW) that was either anaphoric and already given in the context sentence (‘old’) or non-anaphoric (‘new’). The described event or action involving the critical proper name was either coherent or incoherent with the characteristic information described in the context. For example, one target sentence stated that John was the top scorer, which would be coherent with a context that describes John as one of the best players in the team, incoherent with John as one of the worst players, coherent with other names as the worst players, and incoherent with other names as the best players. In other words, we manipulated the factors repetition (old, new) and coherence (coherent, incoherent) in a within-subjects two-by-two design, rendering the critical proper name in the second sentence old and coherent, old and incoherent, new and coherent, or new and incoherent. A coherence rating test described later in this section supports this classification of conditions.

In the fifth, new-neutral condition, the first sentence did not contain proper names but contained the reference group (e.g., *the players in the football team*) as subject of the sentence. In this condition, the critical proper name in the second sentence was new, and it was neutral with respect to the discourse because the context sentence lacked characteristic information about individual referents.

Approximately half (113) of the items only contained typically male names, the other half (112) only contained typically female names. Each name was used in only one item, and each item contained three proper names, meaning that in total 675 names were used. To control for potential effects of order of mention, the old names referred to the first name in the context sentence in one half of the items, and to the second name in the context sentence in the other half. To avoid sentence-final wrap-up effects contaminating the brain’s response in the time window of interest, the CW was always followed by five words that ended the sentence without further anaphoric expressions.

### Filler items

We added 25 coherent filler items without any proper names. The first sentence of the filler items was similar in form to the first sentence of the new-neutral condition, containing a reference group (e.g., *the candidates in the elections*) but no specific individuals. The second sentence contained a definite noun phrase referring to a specific person (e.g., *the politician*) that could belong to the reference group. A translated example of a filler item: *The candidates in the elections are very popular. The majority of the people voted for the politician with the extraordinarily creative ideas*.

### Comprehension questions

We included 80 comprehension questions (50% with correct answer ‘yes’, 50% with correct answer ‘no’) to ensure that participants paid attention to the materials. These had to be answered by means of a button press. Questions were either about the general gist of the mini story or about specific entities in the stories. The average percentage of correctly answered questions was 92.4%. None of the participants scored below the preregistered cut-off of 70%.

### Experimental lists

We created 5 lists, each containing one condition of an experimental item and the same number of items of each condition. Participants therefore only saw one condition of each item. For each of the 5 lists, we created 2 versions by pseudorandomizing the trial order, such that the same condition was never presented more than three times in a row. Each of the 10 lists was presented to 4 participants.

### Coherence rating test

In order to establish that the ‘coherent’ and ‘incoherent’ conditions were indeed considered as such, we conducted a coherence rating test with an additional group of 20 native speakers of Dutch (19 females, average age: 24 years, age range: 18-30 years). In an online experiment, which was done after we had conducted the EEG experiment, these participants rated the coherence of each mini story on a 5-point scale, ranging from ‘not coherent’ to ‘coherent’ (for a similar approach, see Yang, Zhang, Yang, & Lin, 2018). The two sentences of each mini story were presented together. The second sentence was presented up to the critical proper name, allowing us to compare the rating scores to the CW-locked EEG patterns.

Following a recent suggestion by Bürkner and Vuorre (2019), we analyzed the ordinal data of the rating test with Bayesian ordinal regression analysis using the brms (Bayesian regression models using ‘Stan’; Stan Development Team, 2018; Bürkner, 2017, 2018) package for R (version 3.5.0; R Core Team). Coherence (coherent, incoherent) and repetition (new, old) were deviation coded. Both were added as fixed effects, together with their interaction. We included participant and item as random effects, each with coherence and repetition as random slopes. The violin plot of the results, presented in Figure 1, seems to indicate that the variance of the ratings varies across the four relevant conditions. We therefore incorporated unequal variances in the model by adding the variance component of the variables as additional to-be-estimated parameter (Bürkner & Vuorre, 2019).

**Figure 1.**
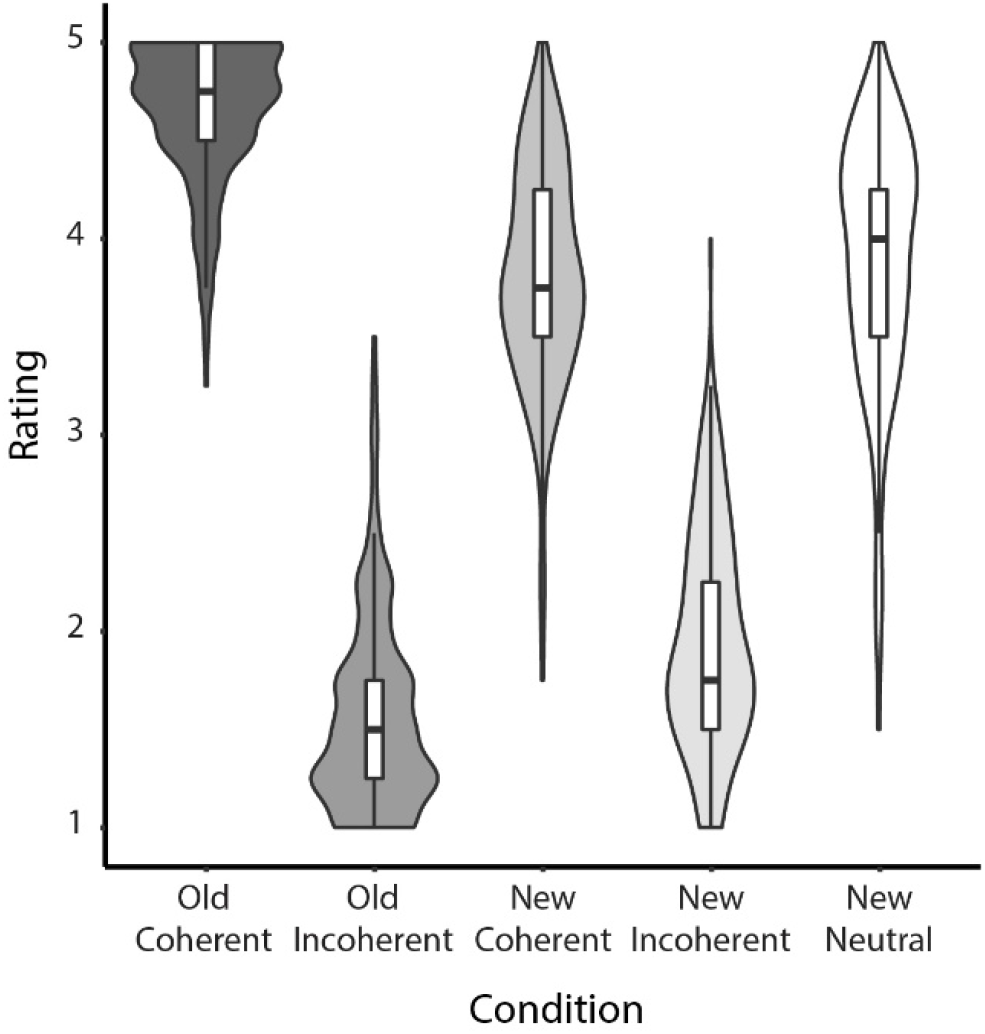
Violin and boxplots of the rating scores per condition, representing the distribution of the average ratings of each item in all conditions.

As can be seen in Figure 1, incoherent items (Mean = 1.75, *SE* = 0.03) were, on average, judged less coherent than coherent items (Mean = 4.24, *SE* = 0.03), *β* = −3.20, CrI = [-3.88, − 2.54]. Old items (Mean = 3.11, *SE* = 0.06) were judged more coherent than new items (Mean = 2.88, *SE* = 0.05), *β* = 0.41, CrI = [0.18, 0.64]. In addition, the difference between coherent and incoherent items was bigger for old items than for new items (mean difference = 1.22), *β* = − 2.35, CrI = [-2.57, −2.15]. The estimates for the variance component revealed that the variance of the ratings of incoherent items was larger than that of coherent items, *β* = 0.85, CrI = [0.76, 0.95]. Moreover, the variance of the ratings of new items was larger than that of old items, *β* = 1.48, CrI = [1.35, 1.63].

These results show that participants noticed the difference in coherence between coherent and incoherent versions of the same item, and that EEG modulations as a function of this manipulation might indeed be associated with a perceived difference in coherence.

### Procedure

Participants were individually tested in a soundproof booth. They were instructed to attentively and silently read sentences for comprehension and answer the comprehension questions. All of the stimuli were presented visually in black letters (font Times New Roman, size 34) on the center of the screen, which had a light grey background.

Each trial started with a fixation cross. When participants pressed a button, the first sentence of each item would be presented as a whole. After they had read the sentence and pressed a button, the second sentence was presented word by word. Each word was presented for 400 ms, with an inter-stimulus-interval of 200 ms. The sentence-final word was presented for 800 ms and was either followed by a fixation cross, indicating the start of the next trial, or by a comprehension question. Participants were asked to minimize eye blinks and body movements during the word-by-word presentation of the second sentence.

The 250 items were presented in five blocks of 50 items. Each block contained 9 items of each condition and 5 filler items. 16 items per block were followed by a comprehension question. Participants were allowed to take short breaks between blocks. In total, the experiment lasted approximately 70 minutes.

### EEG recording

The EEG was recorded using an MPI custom actiCAP 64-electrode montage (Brain Products, Munich, Germany), of which 59 electrodes were mounted in the electrode cap (see Figure 2). Horizontal eye movements (horizontal EOG) were recorded by one electrode which was placed on the outer canthus of the right eye, and eye blinks (vertical EOG) were recorded by two electrodes placed below both eyes. One electrode was placed on the right mastoid, the reference electrode was placed on the left mastoid and the ground was placed on the forehead. The EEG signal was amplified through BrainAmp DC amplifiers, referenced online to the left mastoid, sampled at 500 Hz and filtered with a passband of 0.016-249 Hz. Preprocessing was performed in BrainVision Analyzer (version 2.1; Brain Products, Munich, Germany).

**Figure 2.**
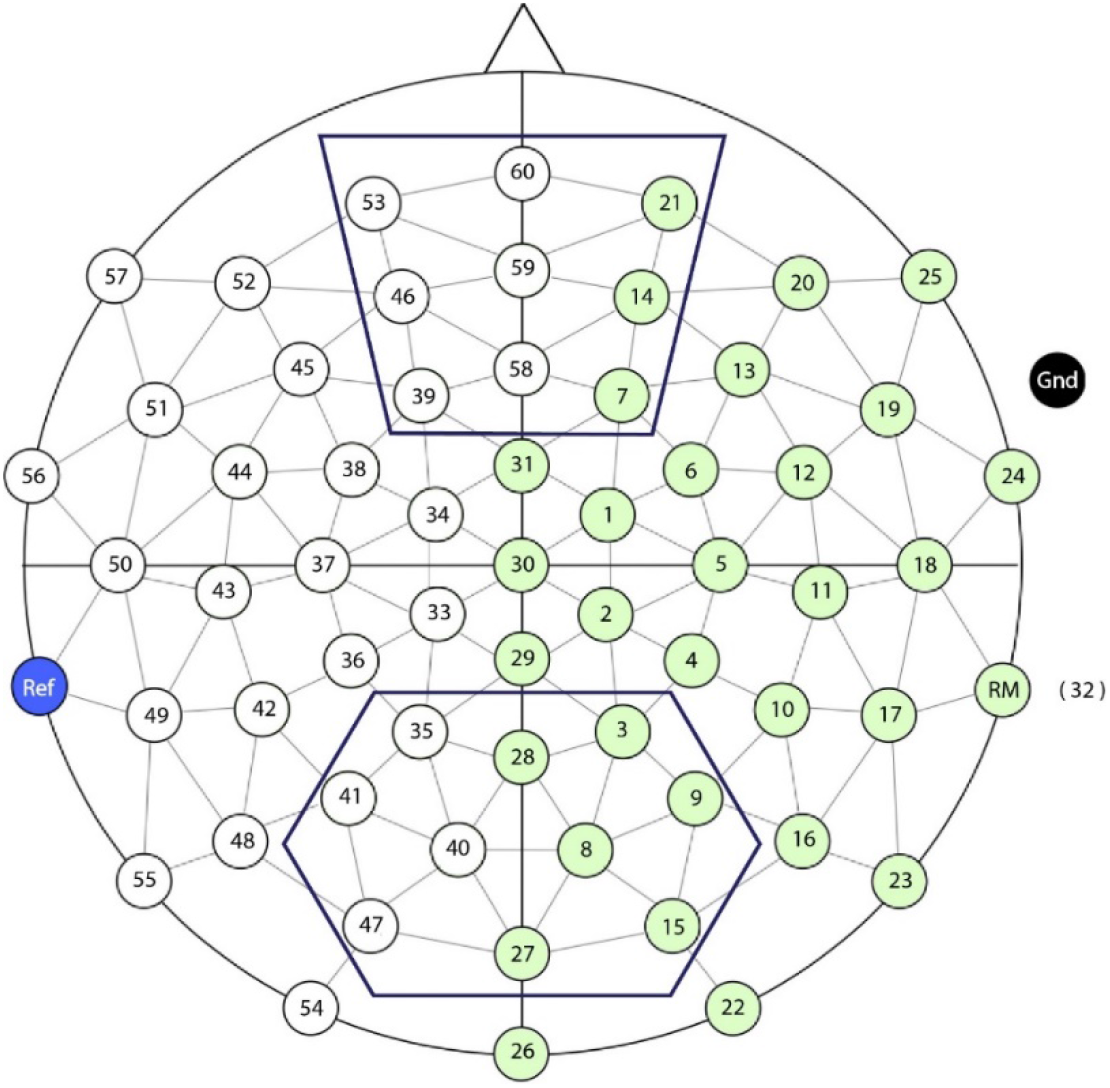
Schematic representation of the 59-electrode array layout. The upper box denotes the Nref region of interest and the lower box the N400/LPC region of interest.

### ERP preprocessing and analysis

We first visually inspected the data and interpolated bad channels if they contained 50 Hz line noise or if they corresponded to broken electrodes. For ERPs, the data was then band-pass filtered at 0.03-40 Hz (24 db/oct) and re-referenced to the average of the left and right mastoids. Segments were created ranging from −500 to 1500 ms relative to CW onset. We visually inspected the data and excluded bad segments containing movement artifacts, multiple-channel muscle activity, or amplifier blocking. Independent Component Analysis (ICA; using ICA weights from a 1 Hz high-pass filtered version of the data) was used to filter artifacts resulting from eye movements and steady muscle activity (Jung et al., 2000). We performed baseline correction using a 250 ms pre-CW baseline interval. Segments containing voltage values exceeding ± 90 μV were automatically rejected. An average of 13.5 segments per participant was rejected (SD = 16.5, average per condition = 2.1). After preprocessing, two participants ended with less than 160 trials and were replaced.

We performed linear mixed-effects analysis (Baayen, Davidson, & Bates, 2008) in R (version 3.5.0; R Core Team), using the lme4-package (Bates, Maechler, & Bolker, 2012). The analyses were done separately for the N400, LPC and Nref regions of interest. At the N400 region of interest, the dependent variable was the average voltage for each trial across the centroparietal electrodes 35, 28, 3, 41, 40, 8, 9, 47, 27, 15 in a 300-500 ms window after CW onset (see Figure 2). At the LPC region of interest, voltage at the same centroparietal electrodes was averaged in a 500-1000 ms window after CW onset. Dependent variables of the N400- and LPC regions of interest were computed separately for each trial and each participant. Coherence and repetition were deviation coded. We started with a full model that included repetition (new, old), coherence (coherence, incoherent) and their interaction as fixed effects, and participant and item as random effects. We also started with a maximal random effects structure by including the interaction term as by-participant and by-item random slope (Barr, Levy, Scheepers, & Tily, 2013). These models did not converge, even after removing the random correlations. We then removed the random slope for the interaction term and retained repetition and coherence as by-participant and by-item random slopes. In order to locate the model with the best fit, we started from the full model and reduced its complexity stepwise, by first removing the interaction and then the main effects. Models were compared using R’s `anova()` function, and *p*-values below *α* = 0.05 were treated as significant.

At the Nref region of interest, the dependent variable was the average voltage for each trial across the frontal electrodes 53, 60, 21, 46, 59, 14, 39, 58, 7 in a 300-1500 ms window after CW onset (see Figure 2). In two separate analyses, we tested the effect of condition, where condition either had the levels ‘new-neutral’ and ‘new-coherent’ or ‘new-neutral’ and ‘old-coherent’. Participant and item were entered as random effects, and condition as by-participant and by-item random slope. We compared the models with and without condition using R’s `anova()` function, treating *p*-values below *α* = 0.05 as significant.

### Oscillatory preprocessing and analysis

For oscillations, the data was band-pass filtered at 0.1-100 Hz (24 db/oct), re-referenced to the average of the left and right mastoids, and segmented into epochs ranging from −1000 to 2500 ms relative to CW onset. We then performed the same procedure for inspection-based artifact rejection and ICA-based correction as for the ERP segments. The resulting dataset for each participant contained many artifact-free trials with voltage values exceeding ± 100 μV. We therefore considered the preregistered ± 100 μV amplitude criterion of the automatic artifact rejection procedure to be too conservative, excluding 22.9 trials per participant (SD = 40.0; where the high standard deviation was driven by one participant). We chose to use a more liberal difference criterion, which excluded segments for which the difference between the maximum and minimum voltage exceeded 200 μV (similar to the difference between −100 μV and 100 μV). This procedure excluded on average 12.2 segments per participant (SD = 23.1; average per condition = 2.3). One participant had fewer than 160 trials and was replaced.

Time-frequency analysis of oscillatory power was performed using the Fieldtrip toolbox (Oostenveld, Fries, Maris, & Schoffelen, 2011), following the same procedure as used by Nieuwland and Martin (2017). In order to find a right balance between time and frequency resolution, we performed time-frequency analysis in two different, but partially overlapping frequency ranges. For the low (2-30 Hz) frequency range, power was extracted from each individual frequency by moving a 400-ms Hanning window with ± 5 Hz spectral smoothing along the time axis in time steps of 10 ms. For the high (25-90 Hz) frequency range, we computed power changes with a multitaper approach (Mitra & Pesaran, 1999) based on discrete prolate spheroidal (Slepian) sequences as tapers, with a 400-ms time-smoothing and a ± 5 Hz spectral-smoothing window, in frequency steps of 2.5 Hz and time steps of 10 ms. On each individual trial, power in the post-CW interval was computed as a relative change from a baseline period ranging from −500 to −250 ms relative to CW onset. Per participant, we computed average power changes for each condition separately.

We used cluster-based random permutation tests (Maris & Oostenveld, 2007) to compare differences in oscillatory power across conditions. This non-parametric statistical test deals with the multiple comparisons problem by statistically evaluating cluster-level activity rather than activity at individual data points, thereby retaining statistical sensitivity while controlling the false alarm rate (Maris, 2012). In brief, the cluster-based permutation test works as follows: first, by means of a two-sided dependent samples t-test we performed the comparisons described below, yielding uncorrected *p*-values. Neighboring data triplets of electrode, time and frequency band which exceeded a critical *α*-level of 0.05 were clustered. Clusters of activity were evaluated by comparing their cluster-level test statistic (sum of individual *t*-values) to a Monte-Carlo permutation distribution that was created by computing the largest cluster-level *t*-value on 1000 permutations of the same dataset. Clusters falling in the highest or lowest 2.5^th^ percentile were considered significant. We used the correct-tail option that corrects *p*-values for doing a two-sided test, which allowed us to evaluate *p*-values at *α* = 0.05.

We compared the items in the old (average of old-coherent and old-incoherent) and new (average of new-coherent and new-incoherent) conditions in the 4-7 Hz theta frequency range in a 0-1000 ms time window and in the 35-45 Hz low gamma frequency range in a 400-600 ms time window. Coherent (average of old-coherent and new-coherent) was compared to incoherent (average of old-incoherent and new-incoherent) in the 60-80 Hz high gamma frequency range in a 500-1000 ms time window. As the cluster-based permutation test is designed to compare two conditions at a time, we tested for an interaction effect by comparing the difference between old-coherent and old-incoherent to the difference between new-coherent and new-incoherent.

#### 2.7.3 Beamformer source localization

In an attempt to identify the sources underlying the observed differences in 4-7 Hz theta power and 60-80 Hz gamma power, we applied a beamformer technique called Dynamical Imaging of Coherent Sources (Gross et al., 2001). This method uses a frequency-domain implementation of a spatial filter to estimate the source strength at a large number of previously computed grid locations in the brain. Because the increase in 4-7 Hz theta activity for old compared to new names was most pronounced between 240-450 ms after CW onset, this time period was subjected to source reconstruction. Following Nieuwland and Martin (2017), the increase in 60-80 Hz gamma activity for coherent compared to incoherent names was analyzed within a 500-1000 ms interval post CW-onset. The procedure and settings of the beamformer approach are adopted from Nieuwland and Martin (2017).

In addition to these condition-specific time windows, we extracted the data of all conditions in a 500-300 ms pre-CW baseline window. These data were re-referenced to the average of all electrodes. For the theta effect we performed time-frequency analysis on 5 Hz, using a Hanning taper with ± 2 Hz spectral smoothing. In the gamma time window, we estimated power at 70 Hz, using a Slepian sequence taper with ± 10 Hz spectral smoothing.

We aligned the electrode positions of the montage to a standard Boundary Element Method head model (a volume conduction model of the head based on an anatomical MRI template; Oostenveld, Praamstra, Stegeman, & Van Oosterom, 2001). This head model was subsequently discretized into a three-dimensional grid with a 5 mm resolution, and for each grid point an estimation of source power was calculated. For the 5 Hz and the 70 Hz frequencies of interest separately, a common inverse filter was computed on the basis of the combined dataset containing the pre-CW and post-CW intervals of both conditions (i.e., old-new for 5 Hz, coherent-incoherent for 70 Hz), which was then separately applied to all trials of each condition in order to estimate source power. After averaging over trials, we computed the difference between post-CW and pre-CW activity for each condition separately in the following way: (post-CW - pre-CW)/pre-CW. In order to visualize the estimated activity, we computed grand averages over participants and subsequently interpolated the grid of the estimated power values to the anatomical MRI.

The estimates of source power were subjected to statistical analysis by means of a cluster-based permutation test. On each source location in the three-dimensional grid we performed a one-sided dependent samples t-test (at *α* = 0.05, yielding uncorrected *p*-values) on trial-averaged data of respectively old and new (5 Hz) and coherent and incoherent (70 Hz). Neighboring grid points with significant *t*-values were clustered. A cluster-level test statistic was calculated by summing the individual *t*-values within each cluster, and evaluated relative to a permutation distribution that was based on 1000 randomizations of the same dataset. In order to localize the spatial coordinates of the areas exhibiting significant differences, we interpolated only the *t*-values of the significant, clustered source points to the anatomical MRI. Brain areas were then evaluated using a template atlas (Tzourio-Mazoyer et al., 2002).

## 3. Results

### ERPs

At the N400 region of interest, the effect of coherence was similar in the old and new conditions (*β* = 0.36, *SE* = 0.38), *χ*^2^ = 0.89, *p* = .35. In addition, coherent and incoherent names elicited similar N400 amplitude (*β* = 0.18, *SE* = 0.27), *χ*^2^ = 0.48, *p* = 0.49. New names elicited a more negative N400 than old names (*β* = −2.14, *SE* = 0.26), *χ*^2^ = 40.75, *p* < .001.

At the LPC region of interest, the effect of coherence was marginally larger in the old than in the new condition (*β* = 0.72, *SE* = 0.39), *χ*^2^ = 3.46, *p* = .06. Coherent and incoherent names elicited similar LPC amplitude (*β* = 0.08, *SE* = 0.28), *χ*^2^ = 0.09, *p* = .76. New names elicited a more positive LPC than old names (*β* = 0.75, *SE* = 0.21), *χ*^2^ = 12.42, *p* <.001. The ERPs and corresponding topographical distributions are depicted in Figure 3.

**Figure 3.**
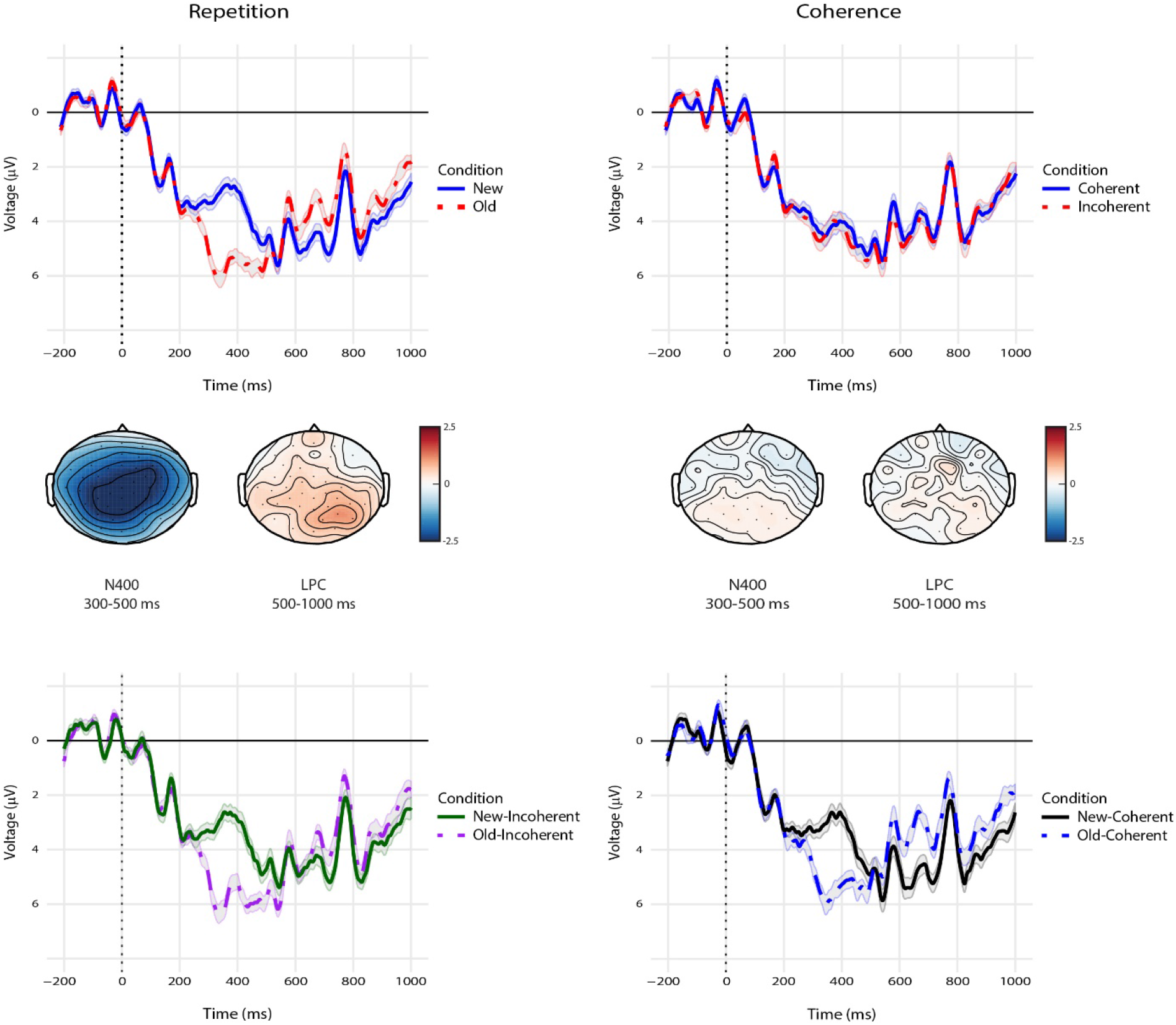
N400 and LPC responses, averaged over the centroparietal region of interest, as a function of repetition (top left) and coherence (top right). Scalp topographies represent the difference between old and new (left) and coherent and incoherent (right) in both time windows. The bottom panel contains the ERPs for individual conditions. In this and all subsequent ERP plots, negative voltage is plotted upwards.

At the Nref region of interest, the average ERP to new-neutral names was more negative than both the average ERP to new-coherent names (*β* = −1.15, *SE* = 0.35), *χ*^2^ = 9.63, *p* = .002, and the average ERP to old-coherent names (*β* = −1.43, *SE* = 0.37), *χ*^2^ = 13.45, *p* < .001. The ERPs and corresponding scalp topographies are shown in Figure 4.

**Figure 4.**
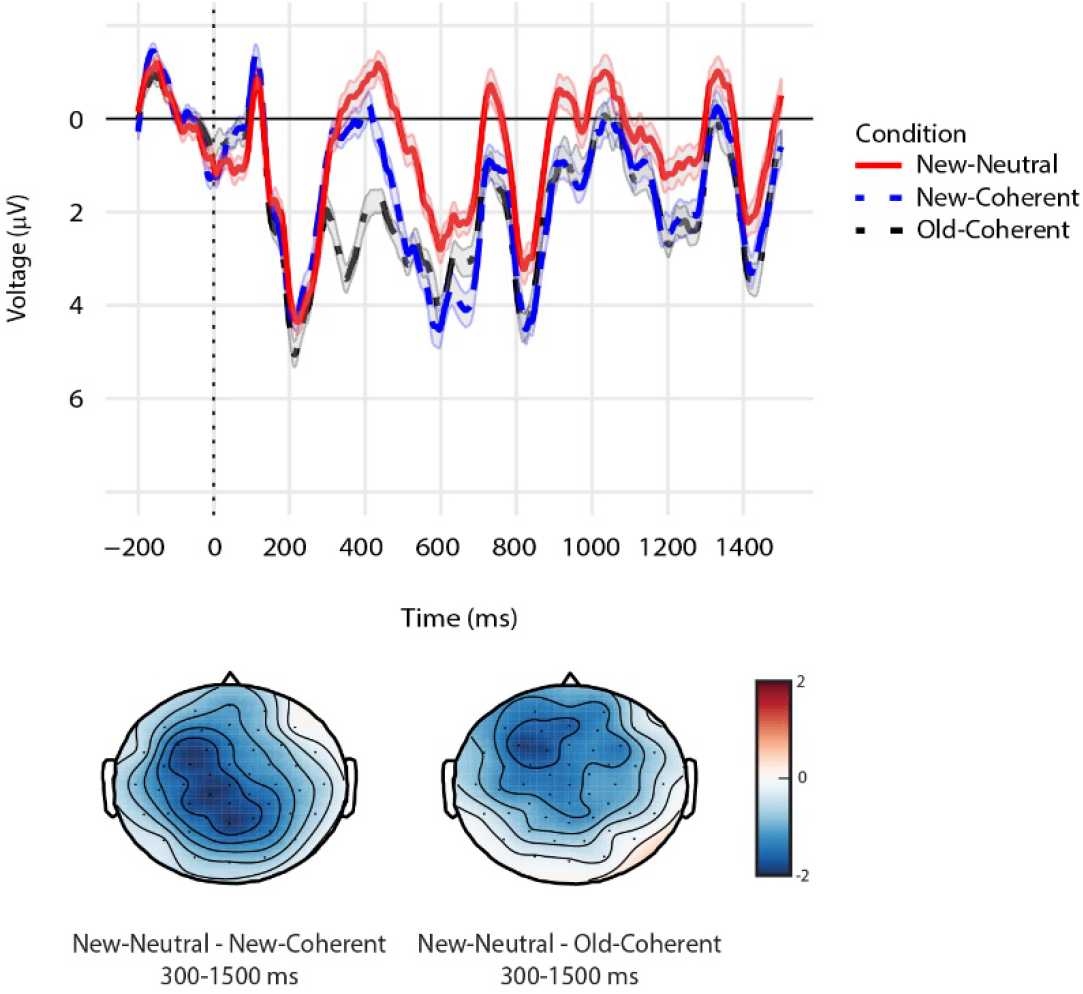
Nref responses, averaged over the frontal region of interest, for the comparisons between new-neutral, new-coherent, and old-coherent. Scalp topographies reflect the difference between new-neutral and old-coherent (left) and new-neutral and new-coherent (right).

### Oscillations

In the 4-7 Hz theta range, we found that old names elicited larger theta power than new names (one significant positive cluster, *p* = .034). This difference was most prominent between roughly 240 and 450 ms after name onset. The time-frequency representations and scalp topography of the theta effect are presented in the left panel of Figure 5. In the 35-45 Hz (low gamma) frequency range, no significant clusters were observed for the contrast between old and new names in the 400-600 ms time window (middle panel of Figure 5). However, the time-frequency representations seem to indicate that old names did elicit enhanced gamma power compared to new names around 400-600 ms, but that the frequency range of this effect is not restricted to 35-45 Hz. To assess this possibility, we exploratorily applied a cluster-based permutation analysis to the activity within a frequency range of 40-55 Hz, in the 400-600 ms time window (see also Van Berkum et al., 2004, who reported a gamma power increase for referentially successful pronouns in this frequency band). This yielded one marginally significant cluster that was larger for old than for new names (*p* = .06). In the 60-80 Hz (high gamma) frequency range, we found that coherent names elicited larger power than incoherent names (one significant positive cluster, *p* = .026) within the time window of 500-1000 ms (right panel of Figure 5). This effect was most prominent between 690 and 1000 ms. No significant clusters were observed for the interaction between coherence and repetition.

**Figure 5.**
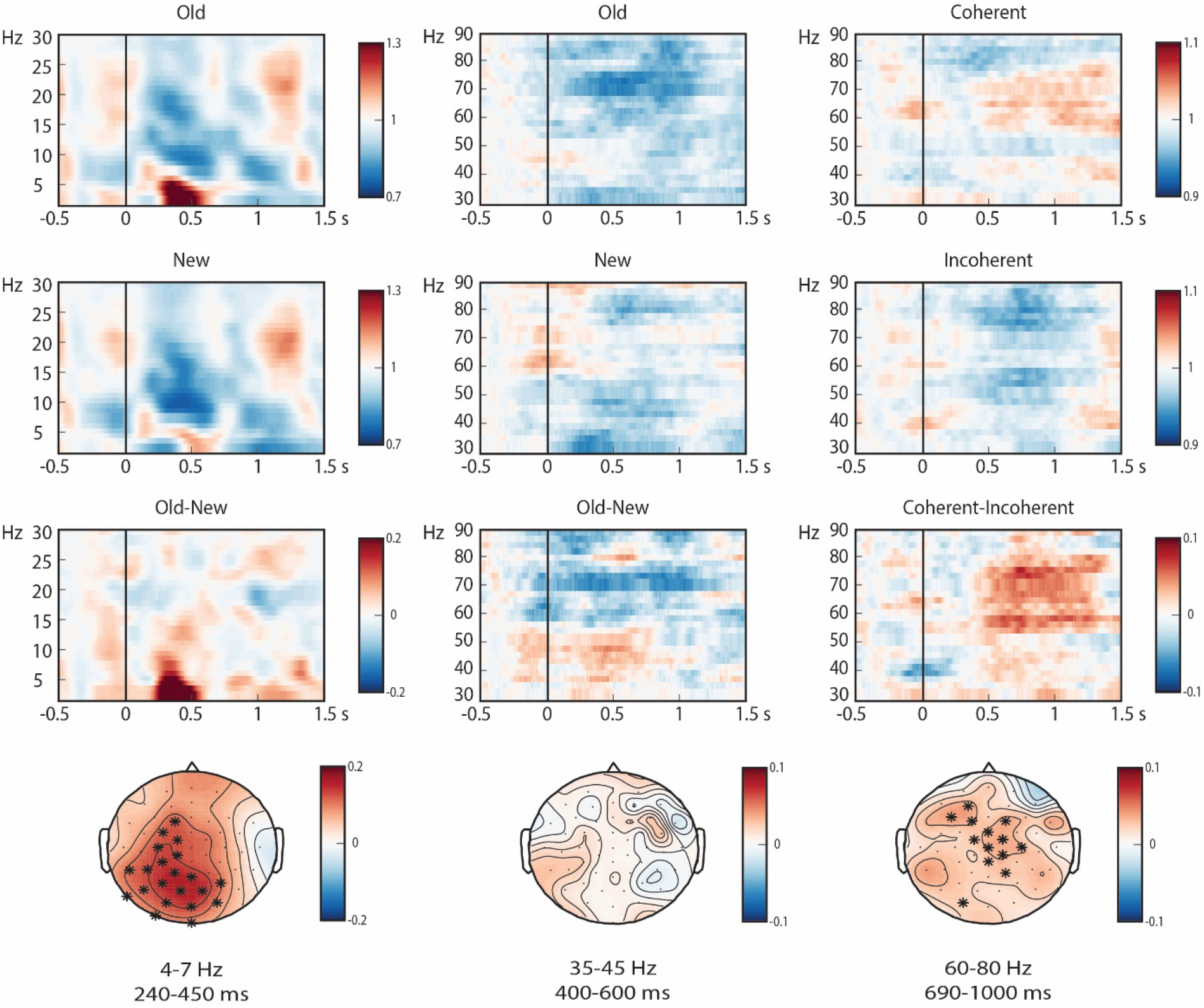
Time-frequency representations and topographical distributions of the theta (parietal-midline electrode 40), low gamma (left parietal electrode 42), and high gamma effects (left frontal electrode 45). The electrodes that showed a significant difference in more than 60% of the selected time windows are marked by * in the topographical plots.

### Beamformer source localization

A beamformer procedure was applied to localize the sources of the 4-7 Hz theta and 60-80 Hz gamma effects. We first applied a spatially unrestricted cluster-based permutation test to the power differences in the entire source space. This did not yield significant sources for either the theta effect or the high gamma effect. We therefore performed exploratory region-of-interest (ROI) analysis of both effects.

While Nieuwland and Martin (2017) did not find effects in the theta range, they anticipated potential involvement of the hippocampus and medial temporal lobe (e.g., Mormann et al., 2005; Nieuwland, Petersson, & Van Berkum 2007). However, restricting the cluster-based permutation test to these areas in the left hemisphere (i.e., hippocampus, parahippocampal region, medial temporal lobe) did not reveal significant differences. Visual inspection of the source power estimations of the theta effect, presented in Figure 6, suggests that the theta effect has a frontal-temporal origin, most prominent in left anterior temporal lobe.

**Figure 6.**
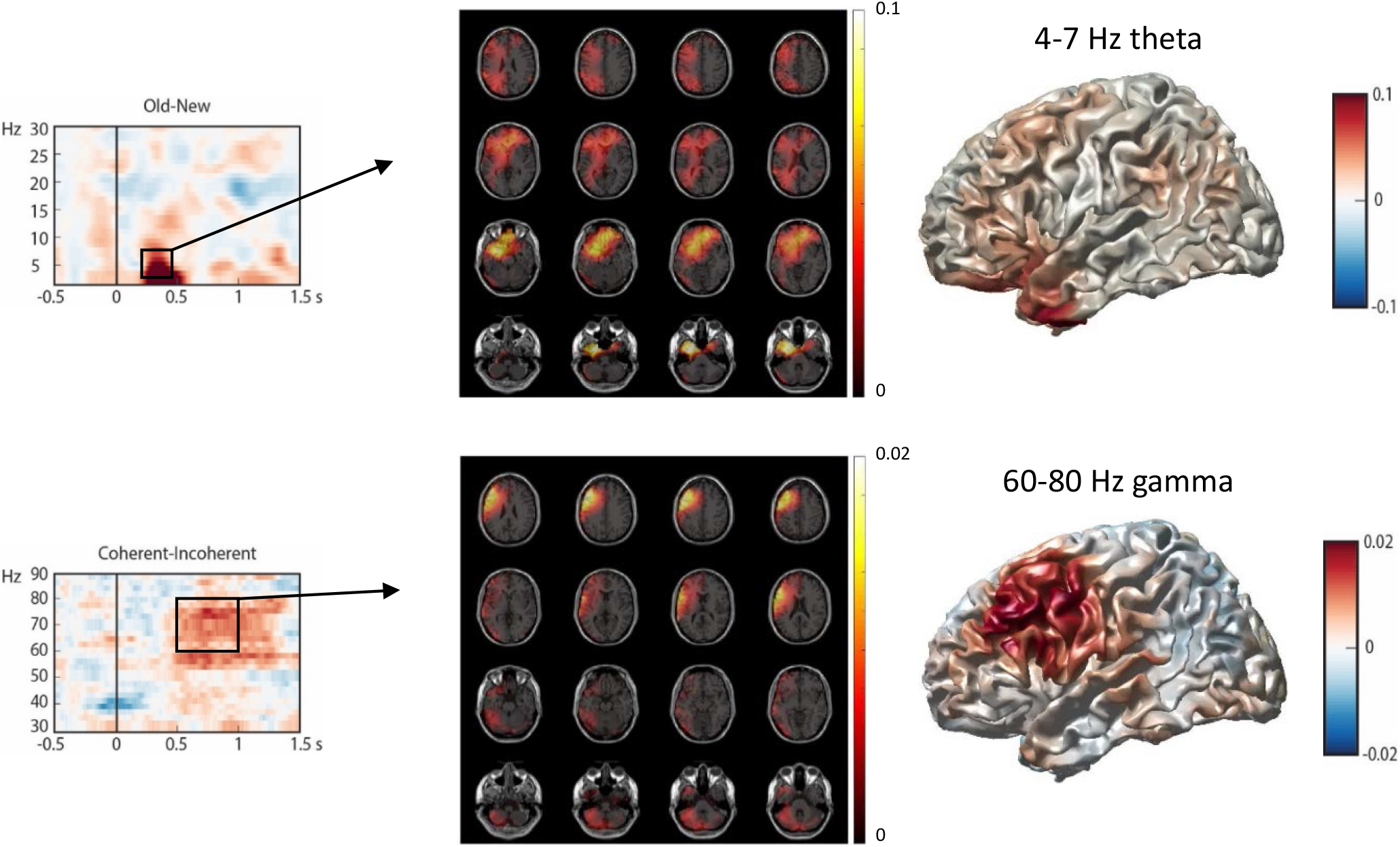
Source localization results for the 4-7 Hz theta effect (top) and the 60-80 Hz gamma effect (bottom). Left: time-frequency representation of the differences, with a black outline indicating the time window of interest. The estimated source power differences are presented in slice view (middle) and surface view (right).

Nieuwland and Martin (2017) localized the source of their high gamma effect to left frontal-temporal regions, encompassing the inferior frontal lobe, inferior temporal lobe and anterior temporal lobe. We performed exploratory ROI analysis by restricting a cluster-based permutation test to these regions. This yielded a marginally significant difference between coherent and incoherent, *p* = .076. Yet, as the effect seems to extend into dorsal regions of the frontal cortex and does not encompass any areas in the temporal lobe (see Figure 6), we performed additional data-driven ROI analysis on the entire left frontal lobe in order to explore where the effect was strongest. In this ROI, a significant difference between conditions was found (one significant positive cluster, *p* = .023), in an area encompassing the left inferior frontal cortex and left medial frontal cortex.

### Exploratory analyses

#### Time-frequency analysis of new-neutral names

The observed old/new theta effect results from an increase in theta power elicited by old names, rather than a decrease in theta power for new names (left panel of Figure 5). This indicates that the effect does not reflect the detection of a mismatch between the new name and the given names mentioned in the discourse context. To provide additional support for this interpretation, we compared theta power elicited by new-neutral names to theta power elicited by both the average of old names (old-coherent and old-incoherent) and the average of new names (new-coherent and new-incoherent). Given that the context sentence in the new-neutral condition does not contain specific referents, these new names should not elicit a mismatch effect. The ‘mismatch account’ predicts a theta effect in the comparison between new-neutral and new names (because the latter might be perceived as mismatching the available referents), but not in the comparison between new-neutral and old names (neither of which should be perceived as mismatching given referents). The theta-band comparison between new-neutral names and both old and new names did not reveal significant differences, neither in a 0-1000 ms time window, nor in a more restricted 240-450 ms time window. Yet, the time-frequency representations depicted in Figure 7 show that, while there is clearly no difference between new-neutral and new names, there seems to be a theta effect in the comparison between new-neutral and old names. This corroborates our conclusion that the theta effect does not reflect a mismatch between a new name and available referents.

**Figure 7.**
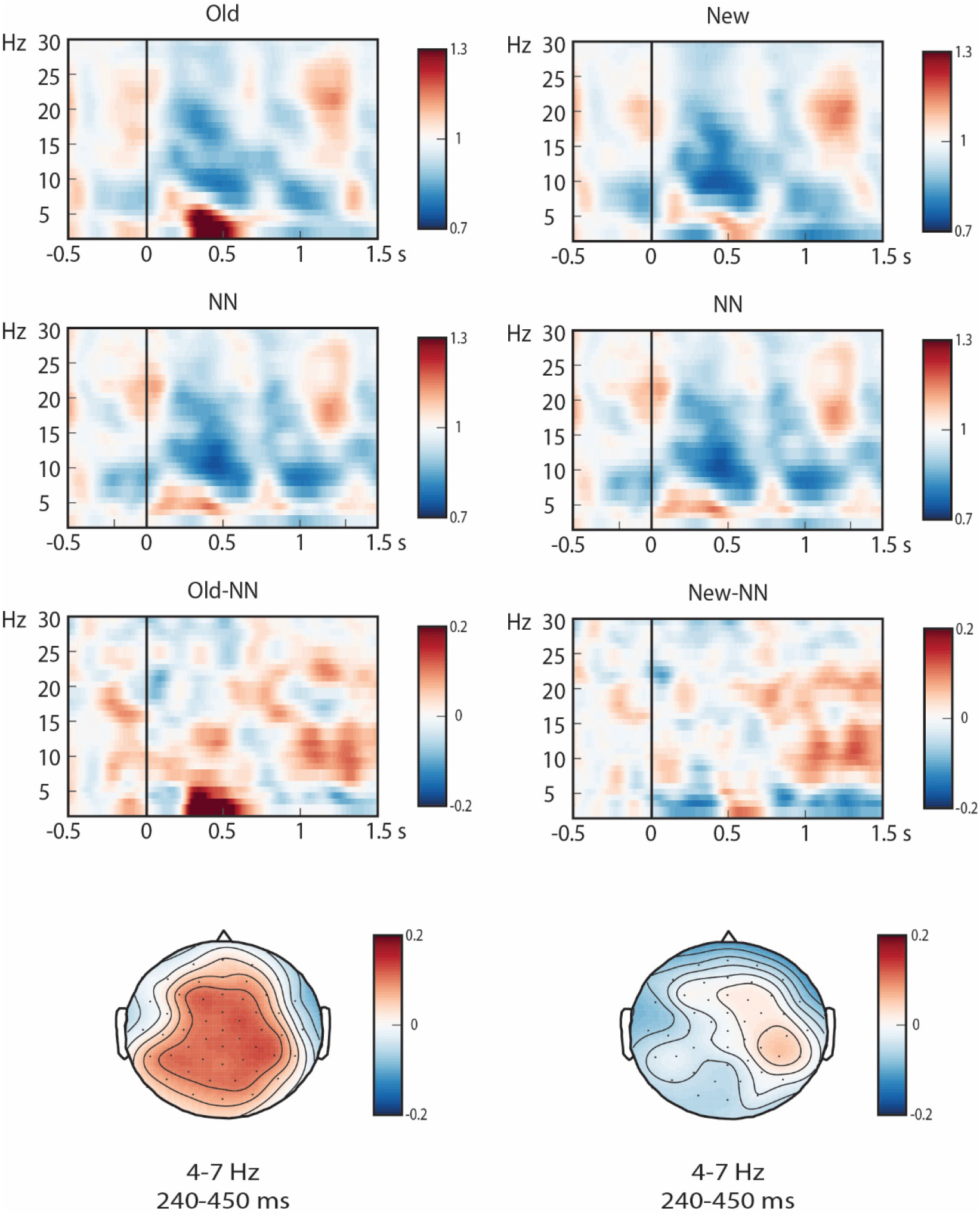
Time-frequency representations and topographical distributions (parietal-midline electrode 40) of the old, new, and new-neutral (NN) conditions. Left: comparison between old and new-neutral. Right: comparison between new and new-neutral.

#### Time-frequency analysis of ERP signal

While ERPs contain phase-locked activity only, time-frequency data contains both phase-locked and non-phase-locked activity. Therefore, the observed differences in the results of the time-frequency analyses might (at least partially) be caused by such phase-locked ‘evoked’ activity. To rule out this possibility, we performed time-frequency analysis on the ERP signal of each condition that was obtained after within-subject averaging (which removes non-phase-locked activity; see Wang, Zhu, & Bastiaansen, 2012 for a similar approach). This analysis was focused on the low frequencies (2-30 Hz; performed in a similar way as described in the section ‘Oscillatory preprocessing and analysis’). A cluster-based permutation test was then used to compare the differences between the conditions old and new within the 4-7 Hz frequency range, both in a 0-1000 ms time window and in a more specific 240-450 time window. Neither analysis yielded significant clusters. Indeed, visual inspection of the right panel of Figure 8 suggests that there is no difference in evoked activity at frequencies above 3 Hz, while our time-frequency analysis of theta oscillations was focused on the 4-7 Hz range. This was shows that the time-frequency results provide information complementary to what can be concluded from the ERP signal.

**Figure 8.**
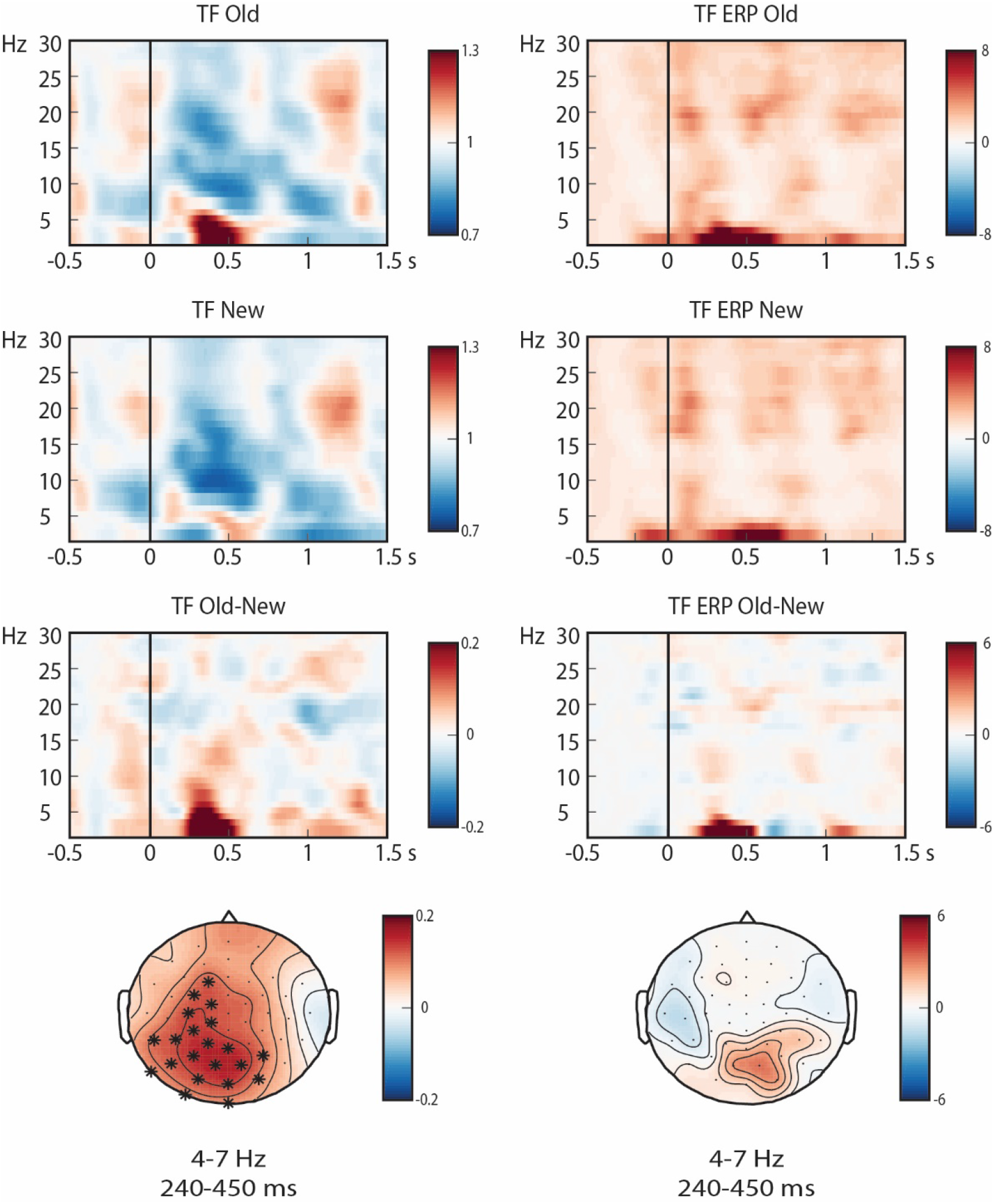
Comparisons of the time-frequency representations (parietal-midline electrode 40) on single trials (left) and on subject-averaged ERP data (right). Topographical distributions of the differences are provided below. The electrodes that showed a significant difference in more than 60% of the selected time windows are marked by *.

## 4. Discussion

This EEG study aimed to tease apart activation and integration of discourse anaphora by analyzing ERPs and neural oscillations. Participants read two-sentence mini discourses in which the interpretation of repeated and new proper names (ease of activation) was either coherent or incoherent with respect to the preceding discourse (ease of integration). As expected, repeated names elicited lower N400 and LPC amplitude than new names. In terms of oscillatory activity, repeated names elicited larger theta (4-7 Hz) power compared to new names. We did not find effects of discourse coherence on either the N400 or the LPC. In the time-frequency domain, however, discourse-coherent names elicited an increase in gamma-band (60-80 Hz) synchronization compared to incoherent names. Beamformer analysis localized this effect to left frontal cortex. We interpret these patterns of theta and gamma synchronization in terms of a two-stage model of anaphor comprehension (Almor & Nair, 2007; Garnham, 2001; Garrod & Sanford, 1994; Gernsbacher, 1994; McKoon & Ratcliff, 1980; Nieuwland & Martin, 2017; Sanford et al., 1983; Sturt, 2003), reflecting respectively activation and integration of discourse anaphora.

### Electrophysiological indices of activation

Repeated names elicited reduced N400 and LPC amplitude compared to new names. We interpret these results to reflect the initial activation of the antecedent (N400) and subsequent updating of the discourse representation by introducing a new referent (LPC). Activation involves the use of relatively superficial information such as lexical or semantic overlap to activate potential antecedent representations in (working) memory (Gerrig & McKoon, 1998; Martin, 2016; McElree, 2000, 2006; McElree et al., 2003). This process is facilitated for repeated names that have just been processed, explaining the reduced N400 (Burkhardt, 2006, 2007; Wang & Yang, 2013; Yang, Perfetti, & Schmalhofer, 2007). The subsequent LPC might reflect updating of the discourse model by establishing an independent referential representation for the new name (Burkhardt, 2006, 2007; Kaan et al., 2007; Schumacher, 2009; Schumacher & Hung, 2012; Wang & Schumacher, 2013). This two-stage interpretation of the N400-LPC complex is largely based on the finding that definite noun phrases (NPs) whose interpretation depends on inferential information (e.g., ‘the bride’ in a story about a wedding) first elicit an N400 that is similar to repeated NPs, and then an LPC similar to new NPs (Anderson & Holcomb, 2005; Burkhardt, 2006, 2007). The reduced N400 suggests that the establishment of a link between two lexically and/or semantically related concepts (*wedding* – *bride*) takes place rather effortlessly, while the LPC effect for new names indicates the cost associated with augmenting the discourse representation with a new, independent discourse entity.

We did not observe a difference between repeated and new names in the preregistered 35-45 Hz (low gamma) range. However, visual inspection of the results suggested that the 35-45 Hz range might have been too narrow, and exploratory analyses suggested that the effect covered a broader frequency range (see also Van Berkum et al., 2004). Although this effect should be interpreted with caution, it is compatible with a role for gamma-band activity in mnemonic processing (see Nyhus & Curran, 2010).

In addition, repeated names elicited an increase in theta-band synchronization, which was most prominent around 240-450 ms after name onset. Time-frequency analysis of the ERP signal in this time window indicated that the effect was not driven by the type of evoked activity that underlies the N400 effect. This corroborates the finding that the old/new ERP and theta effects are complementary phenomena (Chen & Caplan, 2016; Jacobs et al., 2006; Klimesch et al., 2000). In addition, our results suggest that the theta effect does not reflect a mismatch between a new name and specific antecedents available in the discourse context, because it reflected an increase in theta power for repeated names (see left panel of Figure 5) and because it was not elicited by new-neutral names (see Figure 7). We therefore interpret this increase in theta power for repeated referents as reflecting activation of the antecedent in working memory. This interpretation is primarily derived from studies on recognition memory, as recognition memory and anaphor comprehension might recruit similar mnemonic subroutines (Nieuwland & Martin, 2017). Correctly remembered targets elicit an increase in theta-band synchronization compared to correctly rejected distractors (e.g., Burgess & Gruzelier, 1997, 2000; Chen & Caplan, 2016; Jacobs et al., 2006; Klimesch et al., 1997, 2000; Osipova et al., 2006; Van Strien et al., 2005, 2007), which may reflect a relational process that matches the target probe to a representation held in working memory (Chen & Caplan, 2016; Jacobs et al., 2006). Similarly, increased theta power for repeated names might reflect the successful process of matching the repeated name to a representation of its antecedent held in working memory.

This process may rely on feedback projections from the hippocampus to the cortex and top-down control from the (dorsolateral pre)frontal cortex to the hippocampus and posterior cortex (Nyhus & Curran, 2010; Polyn & Kahana, 2008). In line with this idea are the findings of increased theta power at frontal electrodes (Burgess & Gruzelier, 1997; Düzel, Neufang & Heinze, 2005; Klimesch et al., 1997) and increased theta phase synchronization between right frontal and left parietal areas for correctly remembered compared to rejected items (Kim et al., 2012). Source analysis of the theta effect in the current study tentatively suggested that it had a left frontal-temporal origin, including the left anterior temporal lobe (for a discussion of the role of this area in proper name processing, see Semenza, 2011). An important caveat to this discussion is that there are multiple cortical generators of theta oscillations (Raghavachari, Lisman, Tully, Madsen, Bromfield, & Kahana, 2006) and that theta oscillations in different brain regions correlate with task demands (Jacobs et al., 2006). Moreover, given the limited spatial resolution of EEG, the results of EEG source analysis must be interpreted with caution. Future research, using a spatiotemporally more fine-grained methodology such as MEG, needs to shed light on the putative roles of these areas in anaphor comprehension.

### Electrophysiological index of integration

While we did not observe effects of discourse coherence in the ERP analysis, time-frequency analysis revealed an increase in 60-80 Hz gamma-band synchronization for discourse-coherent compared to incoherent proper names. The discrepancy between the outcomes of the ERP and oscillatory analyses might reflect the extent to which their analysis procedures allow variability in the onset of effects. As ERPs are calculated by averaging the EEG signal over a large number of trials, any event-related modulation of the EEG that is not time-locked across multiple trials is strongly reduced in the average ERP signal. If the acquired meaning of the proper names in our experiment was not yet established enough to yield immediate processing difficulty in response to incoherent information, ERPs might not be sensitive to these effects of discourse (in)coherence. Time-frequency analysis, instead, allows for more time variability with respect to the onset of effects (Bertrand & Tallon-Baudry, 2000; Cohen, 2014), and the gamma-band effects seem to corroborate this.

Exploratory region-of-interest analysis localized the gamma-band effect to left frontal regions, encompassing the left inferior frontal cortex and left medial frontal cortex. We interpret this as an effect of semantic integration or unification^8^ (Bastiaansen & Hagoort, 2006; Hagoort et al., 2009). Previous research has related sentence-level unification processes to gamma oscillations, whereby it is generally observed that gamma-band synchronization increases whenever the linguistic input can be integrated into a semantically coherent representation (Bastiaansen & Hagoort, 2015; Fedorenko et al., 2016; Hald, Bastiaansen, & Hagoort, 2006; Peña & Melloni, 2012; Penolazzi, Angrili, & Job, 2009; Rommers, Dijkstra, & Bastiaansen, 2013). The current findings extend this literature by showing that gamma oscillations also index semantic unification on the level of discourse. This further substantiates the idea that information from both sentence-level and discourse-level sources is integrated in a ‘single unification space’, which has been localized to left inferior frontal regions (Hagoort & Indefrey, 2014; Hagoort & Van Berkum, 2007; Van Berkum, Hagoort, & Brown, 1999; Van Berkum, Zwitserlood, et al., 2003). To our knowledge, our results are the first demonstration that gamma-band activity from left frontal cortex tracks the coherence between words and discourse context.

### Nref effect for neutral proper names

Proper names in the new-neutral condition elicited an Nref effect compared to both old-coherent and new-coherent proper names. The Nref has been prominently associated with referential processing (e.g., Nieuwland & Van Berkum, 2008; Van Berkum et al., 2007), and our findings are in line with this view. In the cue-based retrieval architecture (McElree, 2000, 2006; McElree et al., 2003), the second element in a referential dependency (e.g., an anaphor) triggers the activation of already encoded information that is held in working memory (i.e., its antecedent), which is addressable by virtue of content overlap between both representations (Lewis & Vasishth, 2005; Lewis, Vasishth, & Van Dyke, 2006). We suggest that the proper name in the new-neutral condition triggered the activation of the reference group, which can be activated because it overlaps with the proper name on several features (e.g., gender, animacy). However, because the content overlap between anaphor and antecedent is not perfect (e.g., number mismatch), the formation of an anaphoric dependency does not run smoothly, explaining the Nref effect. It is yet unclear whether the Nref effect reflects difficulty of referent activation and/or integration^9^ (Nieuwland & Van Berkum, 2008; but see Karimi et al., 2018).

One caveat to this interpretation is that the new-coherent proper names did not elicit an Nref effect, while this condition also contained a reference group to which the new proper name could be linked (e.g., ‘David and Peter are the worst *players in the football team*i. The top scorer of the team was *John*i). It is possible that the reference group in the context sentence of the new-coherent condition was not accessible enough to be considered available for co-reference. In the new-neutral condition, instead, the reference group was grammatical subject of the context sentence, and this sentence did not contain proper names. Both of these factors have been shown to increase the discourse prominence of the denoted referent (Gordon & Hendrick, 1998; Sanford, Moar, & Garrod, 1988), and might have led to a difference in accessibility of the reference group in the new-neutral vs. the new-coherent condition.

### Implications and challenges for the corticohippocampal theory of reference

Our main results appear to support the corticohippocampal theory of reference (Nieuwland & Martin, 2017), in which anaphor comprehension is a two-stage process that draws upon the brain’s recognition memory network (for referent activation) and the frontal-temporal language network (for referent integration). An important challenge for this theory, however, is specifying how the memory mechanisms underlying recognition memory relate to those underlying anaphor comprehension. Cue-based retrieval models account for the comprehension of anaphora on the basis of the architectural properties of recognition memory. They propose that the activation process underlying linguistic dependency resolution is subserved by a direct-access retrieval mechanism (Martin & McElree, 2011; McElree & Dosher, 1989). In recognition memory studies, direct-access retrieval of working memory representations has been associated with activity in the left inferior frontal gyrus, medial temporal lobe and hippocampus (Nee & Jonides, 2008; Öztekin, Curtis, & McElree, 2008; Öztekin, Davachi, & McElree, 2010; Öztekin, McElree, Staresina, & Davachi, 2008), suggesting a potentially important role for these regions in anaphor comprehension. Consistent with this idea, amnesic patients with hippocampal damage have shown impairments in comprehension of anaphoric pronouns (Kurczek, Brown-Schmidt, & Duff, 2013), and increased activity in the left hippocampus has been observed in response to referentially coherent compared to referentially failing pronouns (Nieuwland, Petersson, & Van Berkum, 2007). Source localization of theta effects in recognition memory studies has also implicated hippocampal, medial temporal and posterior cortical regions (Düzel et al., 2003; Klimesch et al., 2000; Mormann et al., 2005; Osipova et al., 2006). Although these results do not necessarily imply that recognition memory and anaphor comprehension rely on exactly the same neural circuitry, they do suggest the recruitment of similar mnemonic subroutines (Nieuwland & Martin, 2017).

Another challenge for the corticohippocampal theory of reference is explaining how the brain regions for recognition memory and language interact. One possibility, as suggested by Nieuwland and Martin (2017), is that cross-frequency coupling between theta and gamma oscillations allows transient interactions between the relevant neural networks (e.g., Jensen & Colgin, 2007; Lisman & Jensen, 2013). In memory studies, strong correlations between theta phase and gamma power have been observed for successful compared to unsuccessful retrieval (Düzel et al., 2003; Mormann et al., 2005), but theta-gamma coupling has not yet been observed for language-related processes beyond the level of the syllable (Giraud & Poeppel, 2012). It is still an open question whether and how theta and gamma effects in our study are functionally related.

## Conclusion

In this study on the comprehension of proper names in a discourse context, we showed that activation and integration of discourse referents are dissociable in patterns of oscillatory synchronization. Theta (and possibly low gamma) oscillations may play a role in referent activation, and occur within the first hundreds of milliseconds after name onset. We argue that this pattern may be similar to oscillatory effects associated with recognition memory. We extend previous findings by showing that 60-80 Hz gamma oscillations relate to semantic integration on the level of discourse, and occur in a later and more extended time window. Overall, our results suggest a fruitful future for the study of neural oscillations as a potential bridge between the neurobiology of recognition memory and the neurobiology of language.

## Footnotes

1 The distinction between referent activation and integration is similar to that between ‘bonding and resolution’ (Garrod & Sanford, 1994; Garrod & Terras, 2000; Sanford et al., 1983) and ‘recovery and integration’ (McKoon and Ratcliff, 1980).

2 Activation does not have to be fully completed in order for integration to start. Instead, information will be integrated as soon as it is activated, and the resulting discourse representation will be continually updated as new information is activated (see e.g. Rapp & Van den Broek, 2005).

3 The discourse-level N400 effect appears to be identical to the sentence-level N400 effect in terms of latency, morphology and scalp distribution (Nieuwland & Van Berkum, 2006a; Salmon & Pratt, 2002; Van Berkum, Brown, & Hagoort 1999; Van Berkum, Zwitserlood, et al., 2003).

4 Of note, participants were instructed to judge the coherence of context and target on each trial. It is unclear whether such results can also be obtained when no meta-linguistic judgment task is imposed.

5 Notably, integration in this context does not refer to semantic integration, but instead to a process of updating the discourse representation by introducing a new referent.

6 The Nref effect is prominently associated with referential processing (Van Berkum, Koornneef, Otten, & Nieuwland, 2007; Nieuwland & Van Berkum, 2008). Nref effects can be elicited by referentially ambiguous noun phrases (Nieuwland, Otten, & Van Berkum, 2007; Van Berkum, Brown, & Hagoort, 1999; Van Berkum, Brown, Hagoort, & Zwitserlood, 2003) and pronouns (Nieuwland, 2014; Nieuwland & Van Berkum, 2006b; Van Berkum et al., 2004). It has recently been shown that Nref effects can also be elicited by expressions that are not ambiguous but whose antecedents are difficult to activate (Karimi, Swaab, & Ferreira, 2018).

7 After data collection, we found out that for three items the new conditions contained a repeated name instead of a new one. These three items were therefore removed from all further analyses.

8 Some authors argue that these “semantic” gamma-band modulations should be explained in terms of prediction rather than semantic unification (Lewis & Bastiaansen, 2015; Lewis, Wang, & Bastiaansen, 2015; Mamashli, Khan, Obleser, Friederici, & Maess, 2018; Meyer, 2018; Wang, Hagoort, & Jensen, 2018; Wang et al., 2012; following a memory match and utilization model proposed by Hermann, Munk, & Engel, 2004). Indeed, it could be argued that the repeated discourse-coherent proper names in our study were predictable and therefore elicited a gamma-band increase. However, new proper names were never predictable, whether coherent or not, suggesting that some of our conditions do not lend themselves for predictive processes. We therefore do not find the prediction-based explanation of these gamma effects particularly compelling.

9 Another implication of these Nref findings is that proper names are able to trigger antecedent activation (cf. Barkley, Kluender, & Kutas, 2015). Barkley and colleagues (2015) found that pronouns with antecedents elicited a larger anterior negativity than pronouns without antecedents, while no such differences were seen between proper names with and without antecedents. They argued that the proper names did not elicit an Nref because they lack retrieval cues that trigger antecedent activation. Our findings conflict with this claim by showing that proper names do trigger antecedent activation, which allows them to be integrated into a coherent discourse representation.

